# Spatial heterogeneity alters the trade-off between growth and dispersal during a range expansion

**DOI:** 10.1101/2022.04.07.487471

**Authors:** Patrizia Zamberletti, Lionel Roques, Florian Lavigne, Julien Papaïx

**Affiliations:** INRAE, BioSP, 84914, Avignon, France; Université de Rouen Normandie, CNRS, Laboratoire de Mathématiques Raphaël Salem, France

**Keywords:** Eco-evo model, Reaction-diffusion system, Evolutionary R-D trade-off, fragmented space, nonlocal competition

## Abstract

Individuals who invest more in the development of their dispersal-related traits often reduce their investment in reproduction. Thus, there are two possible eco-evolutionary strategies: grow faster or disperse faster (*R* — *D* arbitrage). Here we explore, through a reaction-diffusion model, how spatial heterogeneity can shape the *R* — *D* trade-off by studying the spreading dynamics of a consumer species exploiting a resource in a spatially fragmented environment. Based on numerical simulations and analytical solutions derived from simpler models, we show that the classical mathematical symmetry between the effects of growth and dispersal on the spatial spreading speed is broken in the presence of competition between phenotypes. At the back of the forefront, the dynamics is almost always driven by the *R* specialists. On the forefront, R-strategies are favored in spatially homogeneous environments, but the introduction of heterogeneity leads to a shift towards D-strategies. This effect is even stronger when spatial heterogeneity affects the diffusion term and when spatial fragmentation is lower. Introducing mutations between phenotypes produces an advantage towards the R-strategy and homogenizes the distribution of phenotypes, also leading to more polymorphism on the forefront.

## 1 Introduction

Rapid evolution in species traits can affect their ecological dynamics which in turn feedback on the evolutionary potential (Bonte and Bafort, 2019; Burton et al., 2010). Such interaction between ecological and evolutionary dynamics is crucial to understand the demography when species shift their range as in the case of evolutionary rescue (Anciaux et al., 2019), migrational meltdown (Ronce and Kirkpatrick, 2001), biological invasion (Szűcs et al., 2019). Population expansion is an ecological process mainly driven by traits related to reproduction and dispersal (Deforet et al., 2019; Turchin, 1998). Dispersal affects capabilities to exchange individuals and genes among different habitats (Legrand et al., 2017). Dispersal traits have been proven to be related to body dimension and condition (Duthie et al., 2015; Helms and Kaspari, 2015; Steenman et al., 2015), affecting competitive abilities, food web interactions (Bonte and de la Pena, 2009) or metabolic processes (Hirt et al., 2017). As a consequence, there are many examples where individuals who invest more in the development of their traits related to the dispersal strategy reduce the effort in foraging and reproduction (e.g., reducing their mating period or with lower egg mass) (Baguette and Schtickzelle, 2006; Bonte and Bafort, 2019; Hanski et al., 2006). In such cases, two possible evolutionary strategies exist: dispersing faster or growing stronger (Deforet et al., 2019). This results in a species’ trait trade-off that shapes the ecological and evolutionary dynamics of populations.

During invasion process, there is evidence that trait evolution can be very rapid altering demographic processes (Griette et al., 2015; Perkins et al., 2013). At the forefront, *i.e*., in the frontmost part of the population range, studies highlight that spread rate jointly depends on population growth and dispersal, and that the evolution of these traits can results in an accelerating spread (Fisher, 1937; Perkins et al., 2013). For example, Perkins et al. (2013) focused on how life-history or dispersal traits impact spread rates of the cane toad, *Rhinella marina*, in Australia by combining a stage-structured population dynamics model and an evolutionary quantitative genetic model. They pointed out that rapid evolution of life-history and dispersal traits at the forefront could have led to a more than twofold increase in the distance spread by cane toads across northern Australia. Indeed, spatial sorting of high-dispersal individuals drove dispersal evolution at the forefront and may have resulted in the accumulation of individuals with extreme dispersal abilities at its edge, accelerating invasion (Bouin et al., 2012; Perkins et al., 2013; Shine et al., 2011). However, the individuals leading the forefront should also face novel evolutionary pressures on reproduction, due to low population densities (Kelehear and Shine, 2020). An example of the interactions between dispersal and other key life history traits, such as reproduction, is wing polymorphism of various species of insects (Zera and Denno, 1997). The flight capability (defined by developed wings and flight muscles) is negatively correlated with age at first reproduction and fecundity (Denno, 1994). Thus, the energy efforts for flight and reproduction lead to a trade-off for internal resources (Zera and Denno, 1997).

Reaction-diffusion models are particularly well suited to the study of biological invasions and range expansions in general (Shigesada and Kawasaki, 1997; Turchin, 1998), and to mathematically formalize the relationship between species life-history traits and expansion speed. The first spatio-temporal models of this type considered a homogeneous environment and neglected adaptation (Skellam, 1951). In this case, if the population is initially concentrated in a bounded region, the organisms spread with a speed equal to 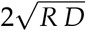 (Fisher, 1937; Kolmogorov et al., 1937), where *R* is the intrinsic growth rate of the population and *D* is the diffusion coefficient which measures the dispersal capacities of the individuals. The population density tends to keep a constant profile: it converges to a traveling wave. The reaction-diffusion framework can be easily adapted to take into account spatial and/or temporal heterogeneities (Shigesada and Kawasaki, 1997). Several theoretical studies considered such models, and proposed a generalization of the notion of traveling wave to spatially-fragmented environments (Berestycki and Hamel, 2002, 2005; Berestycki et al., 2005; Weinberger, 2002). These studies, and other references that we mention in the following sections, have provided a detailed understanding of the dependence of the spreading speed on spatial fragmentation, according to the particular traits they affect (*R*, *D* or both), in the absence of adaptation. In particular, very different effects of fragmentation have been observed, depending on whether they affect *R* or *D* (Hamel et al., 2011).

Some recent works have proposed to take into account genetic adaptation in these spatio-temporal models, thanks to an additional variable, say *y* (interpreted as a phenotypic trait), a mutation term modeled with a Laplace diffusion operator, and a nonlocal selection term (Alfaro et al., 2017, 2013; Alfaro and Peltier, 2021; Peltier, 2020). These models describe adaptation along an environmental gradient, that is, a gradual change in various factors in space that determine the phenotypic traits that are favored by their growth rate *R* (*x, y*). Here, each spatial position *x* is associated with a different optimal trait, *i.e*., a trait which leads to a maximal growth rate. The value of this optimal trait may be proportional to the position (Alfaro et al., 2013; Peltier, 2020), may depend periodically on *x* (Alfaro and Peltier, 2021), or may change with time (Alfaro et al., 2017). Another important part of this literature has been interested in the case where the trait is the diffusion coefficient *D* (Benichou et al., 2012; Berestycki et al., 2015; Bouin and Calvez, 2014; Bouin et al., 2012), and mostly focused on the acceleration of the range expansion in this case, due to the selection of the individuals with enhanced dispersal abilities. The objective of these works was mainly to explain the acceleration of the range expansion of can toads since their introduction in Australia, and the corresponding model is often referred to as the “cane toad equation”. Recently, this framework has been applied to other traits such as the Allee threshold (Alfaro et al., 2021). In all of these reaction-diffusion based models, the additional phenotypic variable only affects a single biological parameter, either directly when this variable is the trait itself such as the diffusion term *D* or the Allee threshold, or indirectly when the growth rate *R*(*x, y*) depends on an abstract trait *y*. This means that trade-off between traits are not considered. Recently, Bouin et al. (2018) considered such a trade-off between dispersal and growth in the cane toad equation. They mainly focused on theoretical mathematical results and on the occurrence of acceleration, in a homogeneous environment, depending on the rate of increase of the mortality term when the diffusion term is increased.

In this work, we develop a reaction-diffusion model to describe the phenotype-spacetime dynamics of a consumer species in a fragmented space during a range expansion. We focus on the trade-off between the growth rate *R*(*x,y*) and dispersal rate *D*(*x,y*), which are both defined as functions of the space variable *x* and the phenotype variable *y*. In a spatially homogeneous environment and in the absence of mutations and Allee effects, the standard formula 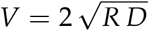 clearly shows that growth and dispersal play a similar role on the spreading speed (Kolmogorov et al., 1937). We analyze here how this symmetry in the effects of *R* and *D* may be broken when facing spatial fragmentation, in the presence of competition between phenotypic traits or in the presence of mutations.

## 2 Model and methods

### 2.1 Eco-evolutionary dynamics

`{model}`

At time *t* and location *x*, the density of the consumer phenotype *y* is defined by *c*(*t, x, y*). We describe the spatial dispersion in a one-dimensional environment with a Laplace diffusion operator, corresponding to random walk movements of the individuals, with a mobility parameter (also called diffusion coefficient) *D*(*x, y*) (Shigesada and Kawasaki, 1997; Turchin, 1998). We assume a one-dimensional phenotype *y* ∈ (*y*_min_,*y*_max_). The mutations between phenotypes are also described with a Laplace diffusion approximation (Hamel et al., 2020; Tsimring et al., 1996) with constant mutation coefficient *μ* ≥ 0. The mutation coefficient *μ* is proportional to the mutation rate (per individual per generation) and to the average mutation effect on phenotype (Hamel et al., 2020). Finally, the population grows logistically with a spatially variable growth rate *R*(*x, y*). Competition occurs locally on the geographical space but globally over phenotypes though a nonlocal term, and is modulated by a parameter *γ*. This leads to the following reaction-diffusion model for the phenotype-space-time dynamics of the consumer population:

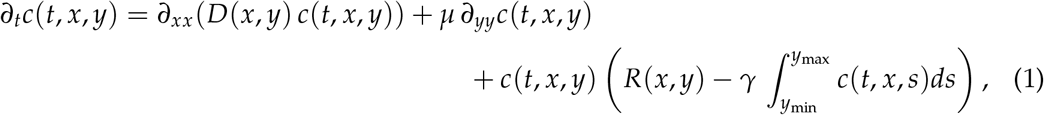

with *t* > 0, 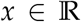 and 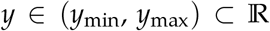. In all cases, we assume an initial condition 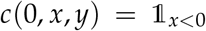, the characteristic function of the domain (*x,y*) ∈ (–∞,0) × (*y*_min_, *y*_max_), and we focus on the spreading of the solution to the right, that is in the direction of positive *x*. In addition, we assume no-flux boundary conditions at the boundaries *y* = *y*_min_ and *y* = *y*_max_:

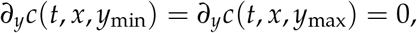

so that in the absence of demography (*i.e*., if *R* = *γ* = 0), and with an integrable initial condition *c*(0,*x,y*), the global population size 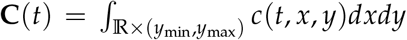 would remain constant.

### 2.2 Modeling genetic and spatial fragmentation in dispersal and growth

Spatial fragmentation in environmental conditions are assumed to impact the consumer growth rate *R* and its mobility *D*. Genetic and environmental effects on *R* and *D* are assumed to be additive. The parameters

*R*_0_ > 0 and *D*_0_ > 0 are the basal values for growth and diffusion. These basal values are modified according to a genetic effect, *R_g_*, respectively *D_g_*, and a environmental effect, *R_s_*, respectively *D_s_*. The parameter *L* controls the spatial fragmentation: *R_s_* (*x*/ *L*) and *D_s_* (*x/L*) are *L*-periodic. A small value of *L* corresponds to a highly fragmented (or rapidly varying) environment, and a large value corresponds to a low fragmented (or slowly varying) environment. Moreover, we also test the effect of introducing an amplitude effect to scale the fragmentation with respect to the scenario presented (see Appendix A).

#### Genetic effect

Given its trait value *y* the genetic effect on the growth rate *R* and the diffusion coefficient *D* is assumed to be Gaussian (Figure 1):

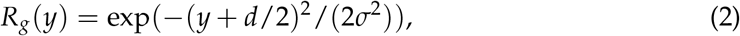

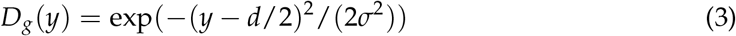

where *d* > 0 corresponds to the distance between the two optima. The optimum trait for diffusion represents the consumer optimal dispersal strategy, and the optimum trait for the growth rate represents the consumer optimal resource exploitation strategy. Here, we assume that the optimum traits are symmetric with respect to 0, *O_R_* = ‒*d*/2 and *O_D_* = +*d*/2 for the growth rate and dispersal, respectively. The coefficient *σ*, fixed to 1 in the following, is the standard deviation of the Gaussian function and indicates the intensity of selection around the optimal trait value (smaller *σ* means higher intensity of selection).

**Figure 1:**
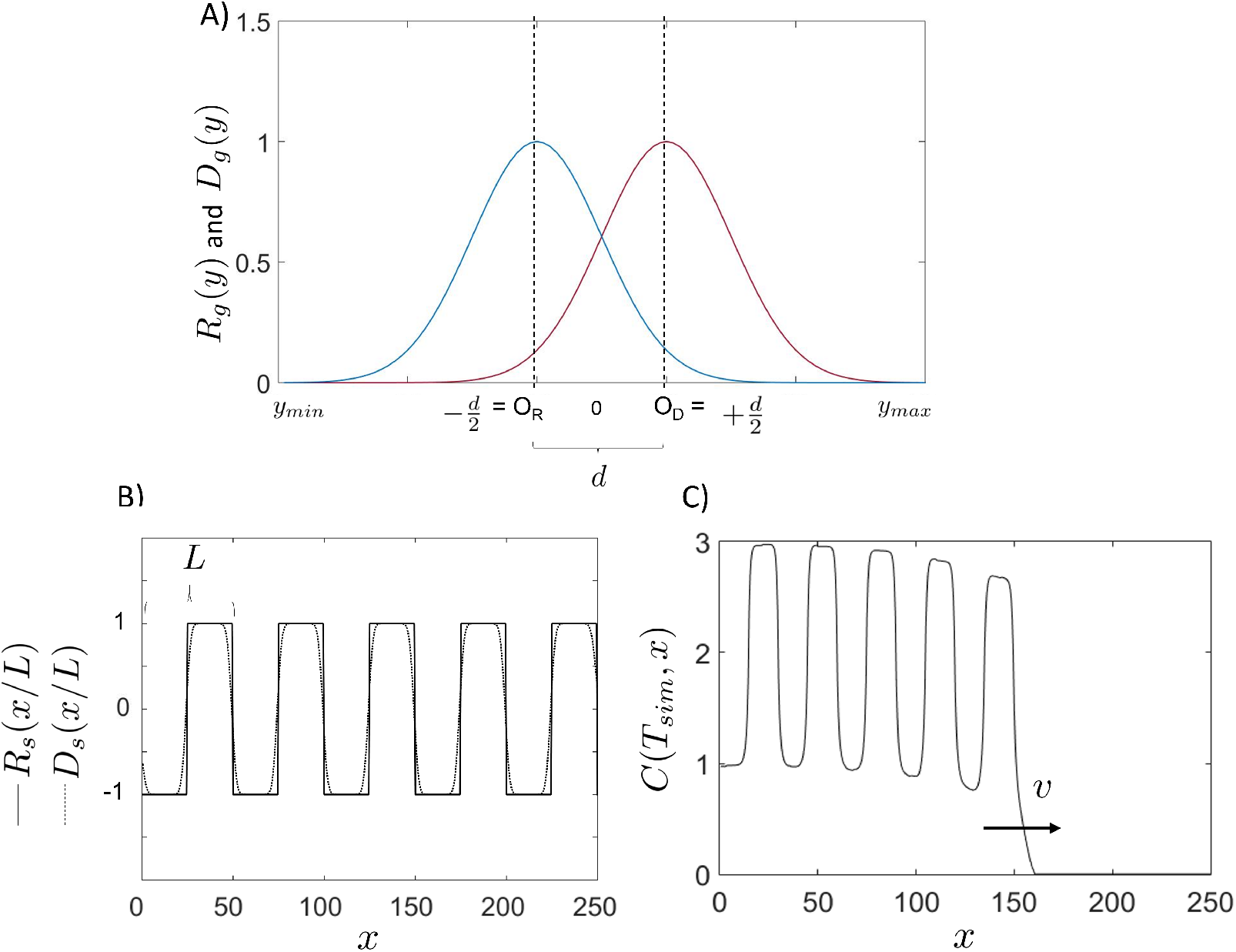
Genetic and environmental effects on growth (*R*) and dispersal (*D*). The panel A displays the curves representing the genetic effect for the dispersal rate *D_g_*(*y*) (red) and growth rate *R_g_* (*y*) (blue) expressed as a function of phenotypic traits *y* ∈ (*y_min_*, *y_max_*). The coefficient *d* is the distance between the optima for dispersal and growth rate. The panel B shows the environmental effect for the dispersal (*D_s_*(*x/L*))(dotted line) and (*R_s_*(*x/L*)) (solid line). The panel C shows an example of resulting total population density *C* (*T_sim_, x*) (see Equation (5)) obtained from the solution of the Equation (1) at time *T_sim_* = 60 along with the position x, with the parameter values: *μ* = 0, *d* = 2, *R*_0_ = 1, *D*_0_ = 1, *σ* = 1 for in the Scenario *R_het_*. `{fig:OdOr}`

#### Environmental effect

The terms *R_s_* (*x/L*) and *D_s_* (*x/L*) describe the periodic variations over the space *x* (Figure 1B). Here, *R_s_* is a 1-periodic piecewise constant function of mean 0, with *R_s_* (*x*) = *R*_0_ on [0,1/2) and *R_s_* (*x*) = – *R*_0_ on [1/2,1). Equivalently, *D_s_* is a smooth 1-periodic function, with mean value 0, and bounded from below by – *D*_0_ (so that *D* is always positive). More precisely, we define the 1-periodic function *δ*_1_ (*x*) such that *δ*_1_ (*x*) = *D*_0_ in [0,1/2) and *δ*_1_(*x*) = – *D*_0_ in [1/2,1).

Then, *D_s_* is obtained by regularizing *δ*_1_ with a convolution by a smooth function:

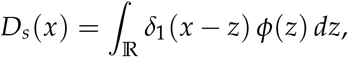

with *ϕ* a Gaussian function with small variance.

### 2.3 Simulation scenario

We define three scenarios, depending on the presence of spatial fragmentation, and on its effects on growth or dispersal:

- Scenario *H*: spatially homogeneous coefficients. In this case,

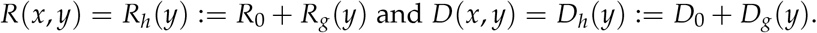
- Scenario *R_het_*: fragmented growth and homogeneous dispersal. In this case,

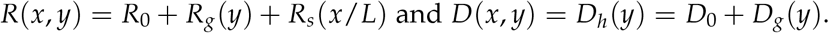
- Scenario *D_het_*: homogeneous growth and fragmented dispersal. In this case,

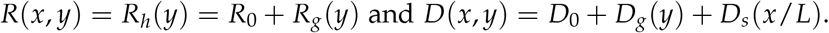

Under these scenarios, we numerically simulated the model of Equation (1), to explore the phenotypic trait composition in the population with different parameter value combinations. Specifically, we focused on: i) the period of fragmentation *L*, considering rapidly (small period *L* = 2) and slowly (large period *L* = 10) varying environments; ii) the distance *d* between the two optima, where we considered a short distance for weak trade-off (*d* = 2) and a large distance for strong trade-off (*d* = 4); iii) and, the mutation parameter *μ* = 0 (no mutations) or *μ* = 0.1. The equations are solved numerically by transforming them into lattice dynamical systems (continuous time, discrete space with small space step), and using a Runge-Kutta method over a fixed spatial domain (defined as *x* ∈ [0;250] by step 0.5). The phenotype space is defined between *y_min_* = —5 and *y_max_* = 5 by step *δ_x_* = 0.1. The implementation is performed by using the software Matlab^®^ (code repository: https://osf.io/v6n4m/).

### 2.4 The spreading speed

`{Speed}`

The consumer species *R* — *D* trade-off is investigated by focusing on the spreading properties and analyzing the population forefront under the defined scenarios. The spreading speed (to the right) *V* is the asymptotic rate at which a population, initially concentrated to the left of some point, expands its spatial range. It can be defined here as the smallest speed such that, if an observer travels to the right *(i.e*., towards increasing *x* values) with this speed, he will observe that the population density vanishes. In mathematical terms, *V* is the only speed such that:

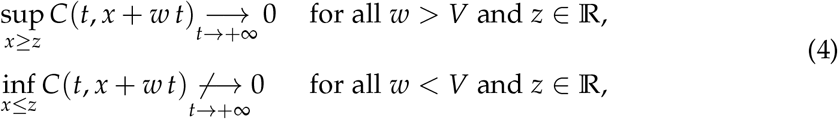

with *C*(*t, x*) the population density at spatial position *x*:

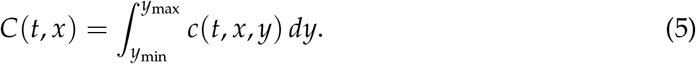

For each phenotype *y*, the spreading speed *v*(*y*) of the phenotype *y* can be defined as well by replacing *C*(*t, x* + *wt*) with *c*(*t, x* + *wt,y*) in the above expressions.

The existence of a spreading speed and analytical characterizations have been obtained for standard equations with spatially homogeneous coefficients and local competition terms (Aronson and Weinberger, 1975, 1978; Fife and McLeod, 1977; Kolmogorov et al., 1937). Comparable results have been obtained with a periodically varying coefficient as in Equation (1) and a local competition term (Berestycki and Hamel, 2002, 2005), namely for equations of the form:

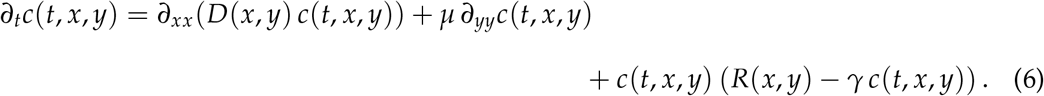

Here, the difference with Equation (1) is that the individuals with phenotype *y* only interact with individuals with the same phenotype. As we did not assume an Allee effect in Equation (1), the solutions should be pulled by the individuals in the leading edge of the colonization (Roques et al., 2012; Stokes, 1976). Their speed should therefore only depend on the growth term through its linearization around 0, here *R*(*x, y*)*c*(*t, x, y*). We therefore conjecture that the spreading speeds *V* of the solutions of the nonlocal Equation (1) and the local equation (6) are equal. This conjecture is supported by the results of Alfaro et al. (2013), which deal with a nonlocal equation of the form (1), with a constant diffusion term *D* and with a growth term of the form *R*(*x,y*) = *r*(*y* — *B x*), with *r*(*y*) = *r*_max_ — *by*^2^ (to each position *x* is attached an optimal phenotype *B x*). This would imply that the fastest phenotype,

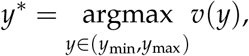

has the same speed for the two Equations (1) and (6) with and without nonlocal interactions. As the fastest phenotype, *y*\* does not compete with other phenotypes, its speed should indeed not be influenced by the competition term, and therefore be the same for the two equations. This conjecture is also supported by the results of Girardin (2017) (Theorems 1.6 and 1.7), who studied an analogue of (1), but with a discrete phenotype space and a spatially homogeneous environment, leading to a system of reactiondiffusion equations coupled by discrete Laplace mutation term.

For Equation (6), under our three scenarios (*H, R_het_, D_het_*), more or less explicit formulas for the spreading speed are available. Thus, we compare these approximations of the spreading speeds with numerical results, using approached models and limiting cases of rapidly and slowly varying environments and we compare these approximations with numerical results.

First, when the environment is spatially homogeneous (scenario *H*), *i.e*., when *R*(*x, y*) = *R_h_* (*y*) and *D*(*x, y*) = *D_h_* (*y*), and in the absence of mutations (*μ* = 0), the spreading speed associated with a phenotype *y* is 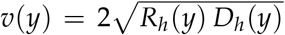 (Kolmogorov et al., 1937). In that case, and according to the values of *d* and *σ* (see Equations (2) and (3)), the fastest phenotypes can be the generalist, *y*\* = 0 or the two specialists, *y*\* = *O_R_* = –*d*/2 and *y*\* = *O_D_* = *d*/2, see Appendix B. The overall spreading speed defined by Equation (4) is 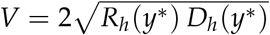. When the environmental fragmentation only impacts the growth rate *R*(*x,y*) keeping the diffusion coefficient spatially homogeneous *D*(*x,y*) = *D_h_*(*y*) (scenario *R_het_*), a general formula for the spreading speed has been obtained by Berestycki and Hamel (2005). Their results also encompass the case of a fragmented diffusion coefficient *D*(*x,y*) but spatially homogeneous growth rate *R*(*x,y*) = *R_h_*(*y*) (scenario *D_het_*). However, in this case, their result holds true for equations with “Fickian” spatial diffusion term, i.e., *∂_x_*(*D*(*x,y*) *∂_x_c*) instead of the Fokker-Planck diffusion *∂_xx_*(*D*(*x,y*) *c*) in (6) (Roques, 2013; Turchin, 1998).

In the spatially fragmented cases (scenarios *R_het_* and *D_het_*) the formulas rely on variational characterizations which make them hardly tractable, even numerically (see Appendix B). More tractable formulas for the phenotype spreading speeds can be obtained for rapidly varying (*i.e*., when the period is small, *L* → 0) and slowly varying (*i.e*., when the period is large, *L* → ∞) environments in the absence of mutations (*i.e*., *μ* = 0) (Hamel et al., 2010, 2011; Smaily et al., 2009). These formulas are summarized in Table 1, see also Appendix B for more mathematical details. We check the accuracy of the analytical approximations in Table 1 and we compare them with numerical simulations for the considered scenarios.

**Table 1:**
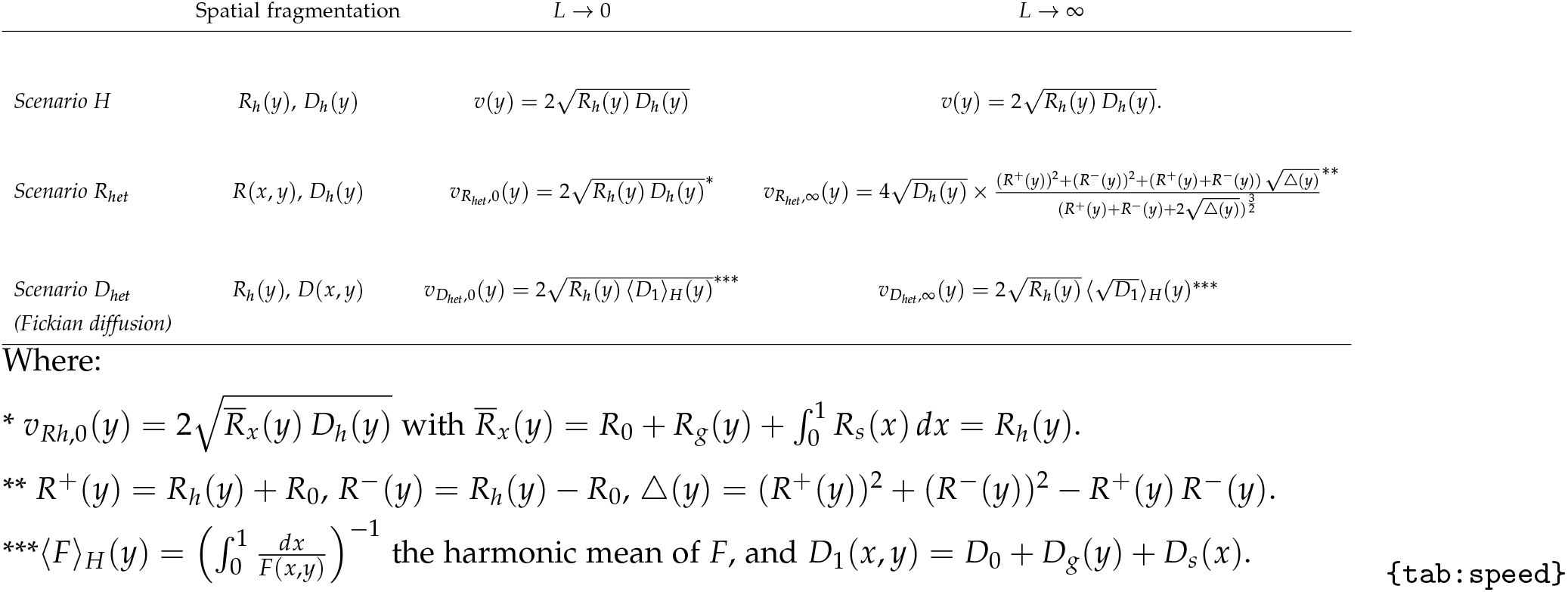
Theoretical phenotype spreading speeds for Equation (6) with *μ* = 0 (no mutation) for rapidly varying environments *L* → 0 and slowly varying environments *L* → ∞.

## 3 Results

`{Res}`

The forefront profiles highlight the role of the *R* – *D* trade-off, the environmental heterogeneity and the mutation in influencing the spreading speed of the different phenotypes (Figures 2–5). In Section 3.1 we analyze these figures, then, in Section 3.2 we use the theoretical formulations to better understand the numerical simulations.

**Figure 2:**
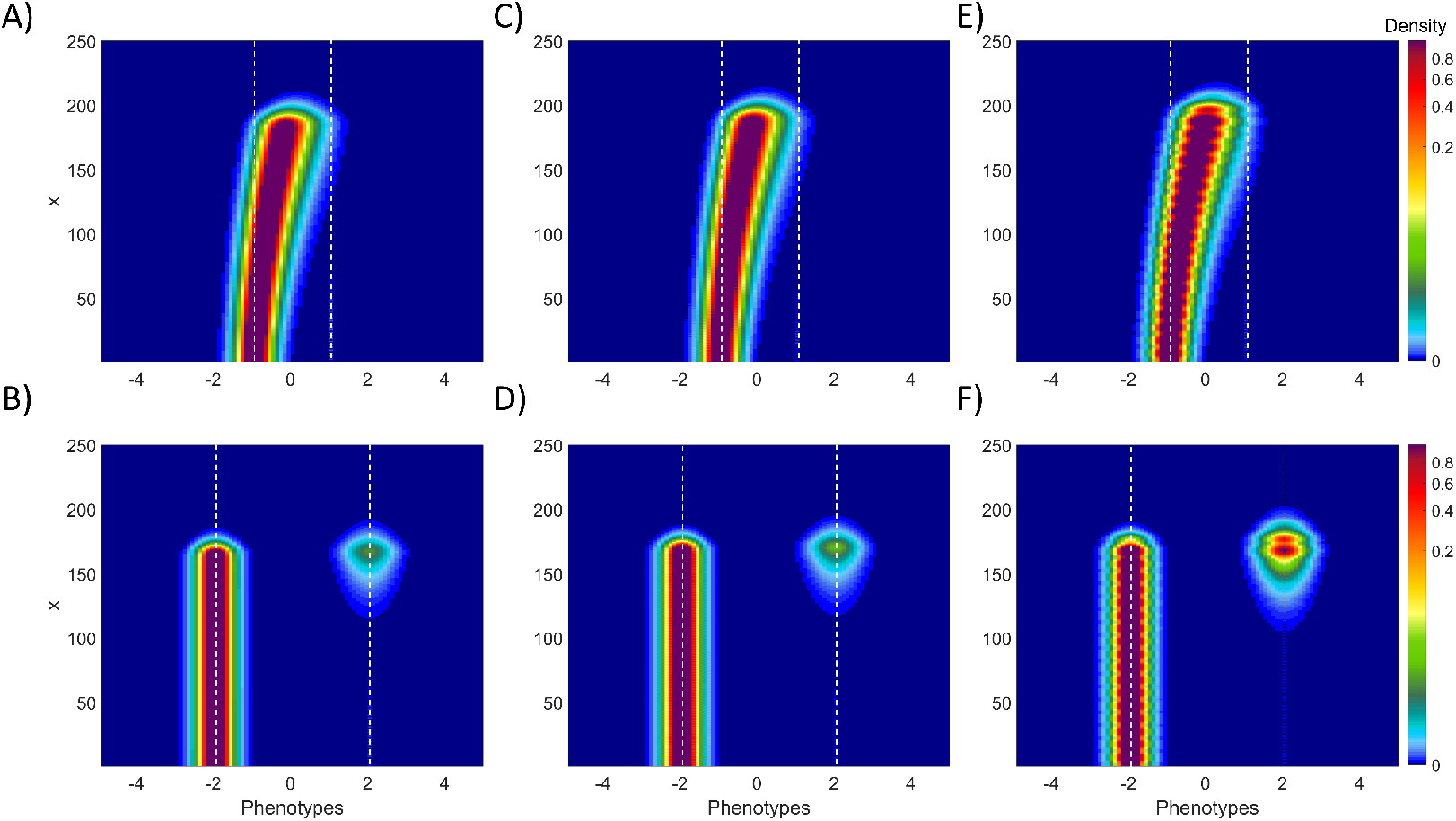
Population density contrasting scenario *R_het_* (Panels C, D, E, F) with scenario *H* (Panels A and B) without mutation. The forefront profiles show the population density with respect to the phenotypes and the space variables. Over lines the tradeoff strength is showed: weak trade-off (*d* = 2) (Panel A, C, E) and strong trade-off (*d* = 4) (Panel B, D, F). Over columns the effect of the period *L* is showed: rapidly varying environment (*L* = 2) (Panel C and D) and slowly varying environment (Panel E and F). Solutions are obtained by numerically simulating the Equation (1) and results are reported at time *T_sim_* = 60. White dashed lines highlight the optimum values (*i.e, O_D_* = *d*/2 and *O_R_* = —*d*/2). `{fig:FrontRh}`

Hereafter, we identify the strategies favoring the selection of phenotypes with a behavior that increases dispersal capacities as D-strategy (namely, when *y*\* > 0), with a generalist behavior as G-strategy (namely, when *y*\* = 0) and with a behavior that increases growth rate as R-strategy (namely, when *y*\* < 0). As at the back of the front the R-strategy is always selected, our focus is entirely dedicated to the trade-off among *R* and *D* on the forefront, see Appendix C.

### 3.1 The R — D trade-off selecting the fastest strategy on the forefront

`{sec:num2D}`

In the absence of mutation (*μ* = 0), when *d* = 2, the forefront is composed mostly by one phenotype with a low degree of specialization on *R* and *D* (Figure 2A, C and E; Figure 4A, C and E). For scenarios *H* and *R_het_*, when fragmentation is high, numerical simulations show a small shift towards the R-strategy (Figure 2A and C, Table 2). Instead, when there is a slowly varying environment, the shift occurs towards D-strategy (Figure 2E, Table 2). Under scenario *D_het_*, whatever environment fragmentation, there is a clear shift towards the D-strategy (Figure 4 C and E, Table 2).

**Figure 3:**
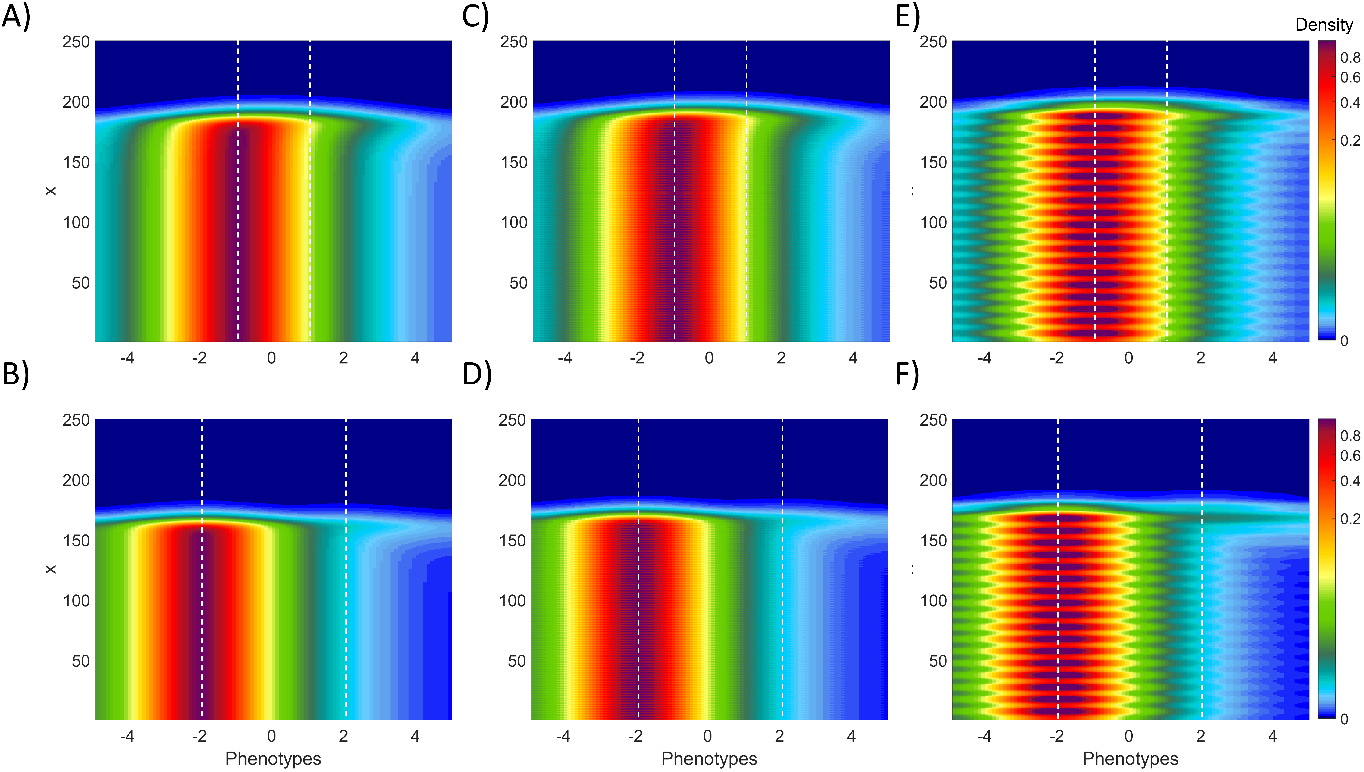
Population density contrasting scenario *R_het_* with scenario *H* with mutation. Caption description is the same of Figure 2, but considering a positive value for mutation (*μ* > 0). `{fig:FrontRhmu}`

**Figure 4:**
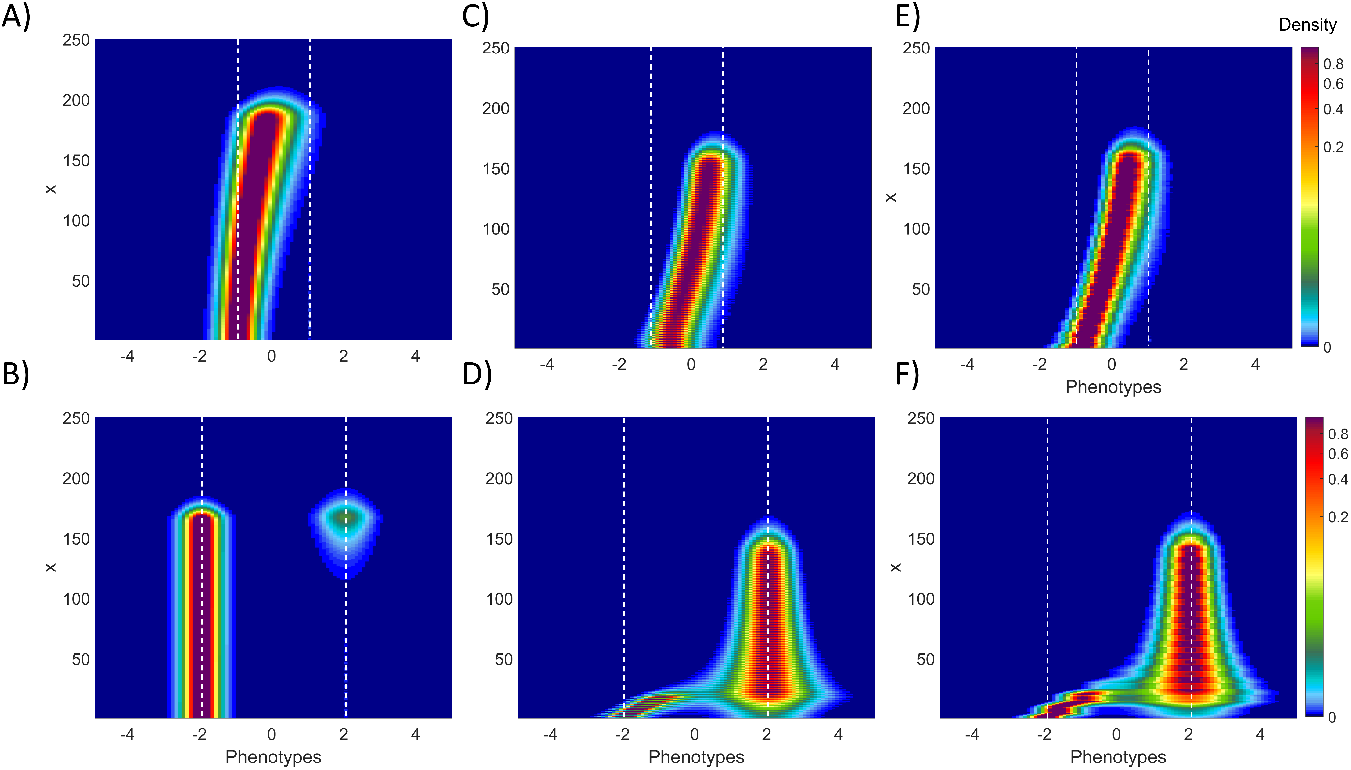
Population density contrasting scenario *D_het_* with scenario *H* without mutation. Caption description is the same of Figure 2, but for scenario *D_het_*. `{fig:FrontDh}`

**Table 2:** Fastest phenotypes leading the forefront. The table reports the values of the fastest phenotype leading the forefront with respect to the optimum value (2*y*\*/*d*). Results of numerical simulations of the Equation 1 are compared over the lines considering the presence and absence of mutation and the theoretical formulas: A) *y*\* = argmax(*v_sim_*(*y*)), with *μ* = 0; B) *y*\* = argmax(*v_sim_*(*y*)), with *μ* > 0; C) *y*\* = argmax(*v_th_* (*y*)), corresponding to the spreading speed *v_th_* reported in Table 1. The corresponding strategy is highlighted: Blue cells correspond to R-strategies (2*y*\* /*d* < 0) and red cells to D-strategies (2*y*\* /*d* > 0). White cells correspond to generalists (2*y*\* /*d* = 0) or when both strategies lead to the same speed (noted R+D). *d* < *d_cr_* correspond to a weak trade-off (*d* = 2), *d* > *d_cr_* correspond to a strong trade-off (*d* = 4). Rapidly varying correspond to a period of heterogeneity *L* = 2 and slowly varying correspond to a period of heterogeneity *L* = 10. `{table:strategi}`

| | | H | $R_{het}$ | | $D_{het}$ | |
| --- | --- | --- | --- | --- | --- | --- |
|  |  |  | Rapidly varying | Slowly varying | Rapidly varying | Slowly varying |
| $d < d_{cr}$ | A | -0.1 | -0.1 | 0.1 | 0.5 | 0.3 |
|  | B | -0.7 | -0.7 | -0.7 | -0.7 | 0.1 |
|  | C | 0 | 0 | 0.2 | 0.6 | 0.5 |
| $d > d_{cr}$ | A | R+D | 0.95 | 1 | 0.95 | 0.95 |
|  | B | -1.2 | -1.3 | -1.35 | 0.5 | 0.8 |
|  | C | R+D | R+D | 1 | 1 | 1 |

When *d* = 4, the colonization is mostly driven by the D-strategy (Figures 2 and 4 B, D and F). A less fragmented habitat (*L* = 10), under the scenario *R_het_*, increases the advantage of the *D*-specialist on the forefront, shifting the trade-off in favour of the D-strategy (see Figure 2 D *vs*. F). For the scenario *D_het_*, the difference between weak and strong trade-off is even more remarkable as the advantage is completely shifted in favor of the strategy *y*\* = *O_D_* (see Figure 4D), defining also a different forefront profile.

These outcomes are completely blurred when introducing a positive value for the mutation coefficient. In fact, the presence of mutations leads to a homogenization of the phenotypic distribution and therefore to a wider phenotype ensemble that leads the forefront: all of the phenotypes should theoretically spread with the same asymptotic speed (see Girardin, 2017, in a homogeneous case with discrete phenotype space). Yet, the population densities (Figures 3 and 5) indicate that the R-strategy becomes the preferred one almost in all cases when *d* < *d_cr_*, except for the scenario *D_het_* with a slowly varying environment (Figure 5 E). The *D*-specialist is still the fastest phenotype under the scenario *D_het_* in case of strong trade-off (*d* = 4) (Figure 5D and F). However, we notice that the shape of the solution at the leading part of the expansion is quite unusual. In all cases, we observe a “bump” corresponding to a fraction of the population which adopts the D-strategy. Thus, the expansion may take advantage of the larger diffusion coefficient of the *D*-specialists and of the larger growth rate of the *R*-specialist by allowing more polymorphism at the leading edge of the propagation.

**Figure 5:**
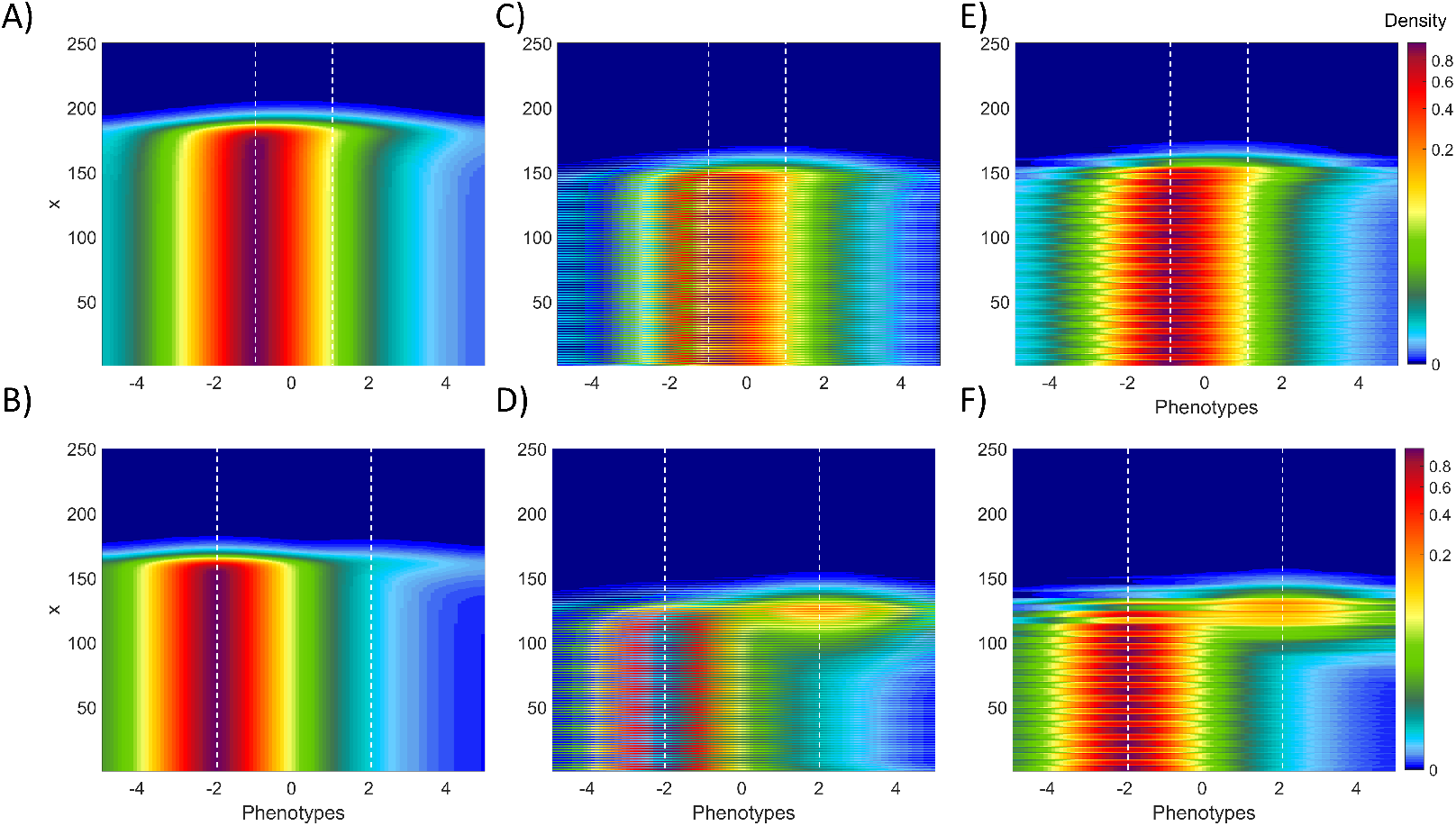
Population density contrasting scenario *D_het_* with scenario *H* with mutation. Caption description is the same of Figure 2, but for scenario *D_het_* with a positive value for mutation *μ* > 0. `{fig:FrontDhmu}`

### 3.2 Insight from the theoretical speeds

`{sec:theor_spee`

In this section, we compare the numerical simulations of Equation (1) presented in Figures 2–5 with the analytical formulations presented in Figure 6 and Table 1. We first check if the outcomes that can be obtained from the theoretical speeds match with the numerical results. These outcomes are summarised in Table 2. Second, when there is a good match, we use the explicit formulas to explain the observed trends. We recall that the theoretical speeds of Table 1 were derived with the simpler local model (6) with *μ* = 0 and therefore do not take the nonlocal competition and mutation effects into account.

**Figure 6:**
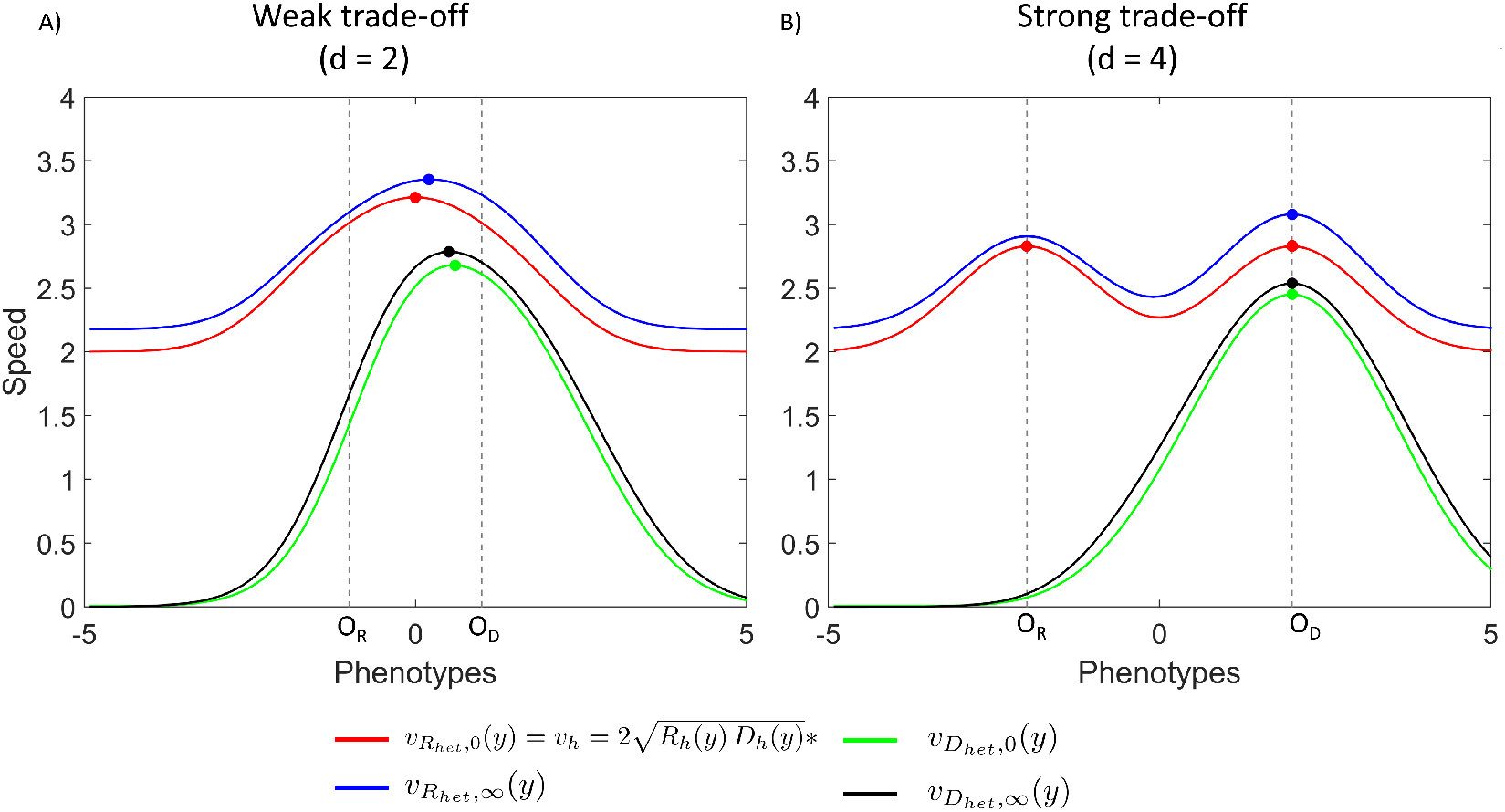
Theoretical phenotype spreading speeds. The theoretical phenotype spreading speeds presented in Table 1 are showed in function of phenotypes *y* ∈ [–5,5] considering a weak trade-off (*d* = 2) (Panel A) and a strong trade-off (*d* = 4) (Panel B). Different colors refers to the formulations of the spreading speed highlighted in Table 1, the dots represent the fastest phenotype leading the forefront. Dashed lines highlight the positions of the optimum traits *O_R_* and *O_D_*. `{fig:v_th}`

In absence of mutation (*μ* = 0), there is a critical threshold *d_cr_* on the distance *d* between the optima (*d_cr_* ≈ 2.4 with our parameter values, see Appendix B), such that the function 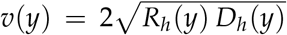, corresponding to the spreading speed, either admits one (*d* < *d_cr_*) or two (*d* > *d_cr_*) maxima. When *d* < *d_cr_* (i.e, *d* = 2) and with a rapidly varying environment, the generalist behavior is expected to be selected as the fastest trait following the theoretical formulations under scenarios *H* and *R_het_*. However, numerical simulations show a small shift towards the R-strategy (Table 2). Instead, when there is a slowly varying environment, or when heterogeneity impacts *D*, theoretical formulations consistently predict a shift to wards the D-strategy (Table 2, Figure 6).

In the scenario *D_het_*, the theoretical speed 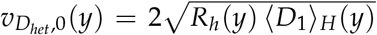 in rapidly varying environments (small period *L* = 2 in the numerical results, *L* → 0 in Table 1) involves the harmonic mean of the diffusion term. Contrarily to the arithmetic mean, the harmonic mean gives a higher weight to small values. Thus, small values of *D*(*x, y*), even on a very small spatial interval, should lead to small speeds *v*_*D*_*het*_,0_(*y*), based on the results of Table 1. This leads to an imbalance in favor of the D-strategies (compare *v*_*D*_*het*_,0_ (*y*) and *v*(*y*) in Figure 6; Figure 4A and B *vs*. Figure 4C and D), which avoid very small values of *D*(*x,y*). In the case of slowly varying environments (large period *L* = 10 in the numerical results, *L* → +∞ in Table 1), the theoretical speed 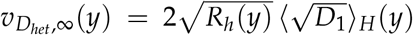 again involves an harmonic mean of the diffusion term (here, its square root) which explains the advantage of the D-strategy, as in the case of rapidly varying environments.

When heterogeneity is introduced on *R* (scenario *R_het_*), in rapidly varying environments, the theoretical formulas predict that the strategy remains unchanged (G when *d* < *d_cr_* or R+D when *d* > *d_cr_*) compared to the homogeneous scenario (H). The theoretical spreading speed of each phenotype in the scenario 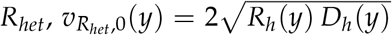, is indeed the same as the speed *v*(*y*) obtained in the homogeneous scenario *H* (both curves are superimposed in Figure 6): as the spatial arithmetic mean of the growth rate (noted 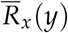 in the legend of Table 1) is precisely equal to *R_h_*, the homogenization results of Smaily et al. (2009) imply that the two speeds are equal. In the numerical simulations (see Table 2, lines A for numerical and C for theoretical), there are some discrepancies, as with a rapidly varying environment (*L* = 2), the homogenization limit is not reached. Hence, we observe a slight shift of the G-strategy towards the R-strategy when *d* < *d_cr_*. When *d* > *d_cr_*, the two strategies are present on the forefront, with a slight advantage for the D-strategy (Figure 2A and B *vs*. Figure 2C and D). In slowly varying environments, the theoretical formulas and the numerical simulations consistently predict a shift towards the D-strategy. In this last case, the formula for *V*_*R*_*het*_,∞_ (*y*) can be written in the form 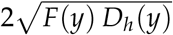, for some function *F* which satisfies *F*(*O_D_*) ≈ (32/27) *R*_0_ (to be compared with *R_h_* (*O_D_*) ≈ *R*_0_) and *F*(*O_R_*) ~ *R*_0_ + 1 for small *R*_0_ (to be compared with *R_h_*(*O_R_*) = *R*_0_ + 1). Thus, compared to the homogeneous case 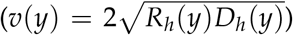, the heterogeneity on *R* creates an asymmetry in favor of the D-strategy which manages to keep a growth rate larger than *R*_0_.

When *μ* > 0 (see Table 2 line B), as expected, there are some discrepancies between the numerical results and the theoretical predictions. However, the arguments above may explain some of the observations. First, in the scenario *D_het_* in most cases (except when *d* < *d_cr_* in rapidly varying environments) we again observe a shift towards D-strategies which is most probably due to the “harmonic mean” effect described above. We note that this shift is stronger in slowly varying environments. In the scenario *R_het_*, although the R-strategy is always selected when *μ* > 0, the positive effect of the heterogeneity on the maintenance of the D-strategy which we noted above in the theoretical formulas is still visible on Figure 3F, which shows more polymorphism compared to the scenario H.

We note the positive effect of increasing the period *L* (equivalently, reducing the environmental fragmentation) on the spreading speeds: this effect, which is obvious in Figure 6 can also be observed in Figures 2–5.

## 4 Discussion

Dispersing faster or growing stronger? In this work, we studied which of these strategies is selected in populations invading a heterogeneous environment. We gathered analytical solutions from the literature and performed numerical simulations of a reaction-diffusion model describing the demo-genetic dynamics of a population invading a onedimensional environment. Results show that the symmetrical effects of growth and dispersal on the spreading speed is broken in the presence of competition between phenotypes, shrinking the population density around the optimum values. From here we observe that, at the back of the forefront, the dynamics is almost always carried out by the *R*-specialists, while, on the forefront, the selection of the fastest strategy is less obvious.

In this study, we identify the main following results: i) R-strategies are favored in spatially homogeneous environments, but the introduction of heterogeneity leads to a shift towards D-strategies, with at least more polymorphism at the forefront; ii) due to a “harmonic mean effect” that we have highlighted through analytical expressions obtained with a simpler model, this phenomenon is even stronger when spatial heterogeneity affects the diffusion term. In this case, the introduction of spatial heterogeneity can lead to a complete switch from an R-strategy to a D-strategy; iii) the spatial fragmentation does not affect a lot the *R* — *D* trade-off, but tends to modulate the polymorphism: in situations where only R-strategists are present at the forefront when the level of fragmentation is high (small *L*), both R-strategists and D-strategists tend to be present at the forefront in low fragmented environments made of large patches (large *L*); iv) mutations produce an advantage towards the R-strategy, and homogenize the phenotype distribution, also leading to more polymorphism on the forefront; v) these effects can be observed with a weak trade-off (such that the generalist *y*\* = 0 leads the population in a homogeneous model without interactions), but become even stronger with a strong trade-off (such that R—strategists and D—strategists have the same speed in a homogeneous model without interactions) and vi) the comparison among theoretical and numerical simulations allows checking when formulas obtained with a simpler model lead to results which are consistent with of a more complex one.

Some of these results (points i,ii,iii) are in accordance with the ones of Burton et al. (2010), who used an individual-based spatial model to study the evolution of three traits in a population undergoing range expansion. When resources are highly fragmented, the trade-off favors to the selection of an R-strategy on the forefront as high resource availability and fecundity facilitate expansion by increasing population growth. By contrast, in a low fragmented environment, the faster dispersers take advantage of their mobility to reach the most favorable habitats and lead the forefront. Evolution thus leads to the selection of a greater capacity for dispersion. Conversely, when heterogeneity impacts dispersal, only the *D*-specialists confer the maximal speed and persist on the forefront, whatever the level of spatial fragmentation. Recently, given two species having growth and dispersal coefficients *R*_1_, *D*_1_ and *R*_2_, *D*_2_ (for species 1 and 2 respectively), Deforet et al. (2019) found that the evolutionary outcome mainly depends on the simple condition 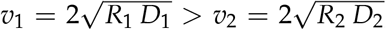, with success of the fastest (here, species 1). In our work, the use of the theoretical formulas of Table 1 mainly relies on the same assumption, that the fastest trait drives the expansion. In most cases, our results show that the observations of Deforet et al. (2019) remain true in a more general context with a continuum of traits and possibly mutations between traits. However, we also observed some discrepancies in the presence of mutations, which tend to advantage the R-strategy. Additionally, as recently observed by Keenan and Cornell (2021) in a homogeneous environment, in the presence of mutations, the furthest forward phenotypes are not necessarily those associated with the largest value of the product *R D* (see below).

Duputié and Massol (2013) argues that natural selection tends to favor dispersal to face spatio-temporal variation in local conditions. Consequently, more dispersive phenotypes are expected to predominate in unstable habitats, while less dispersive phenotypes are common in stable habitats and populations. Here, we found that in the bulk of the population, which corresponds to a saturated population, the R-strategy is always preferred, which is not always the case at the forefront. By definition, the expanding part of the population encounters a more variable environment, especially when the environment is itself highly heterogeneous. In such cases, we observed a shift towards the D-strategy. These findings are also consistent with the “Spatial sorting theory” which predicts that, at the forefront, dispersal may be strongly favored because of the accumulation of the best dispersers (Phillips et al., 2008; Shine et al., 2011; Travis and Dytham, 2002).

The presence of mutations homogenizes the spreading speed between morphs, even in presence of nonlocal competition. We also establish that polymorphism, caused by mutation, is maintained in the presence of spatial fragmentation impacting the *R* and *D* coefficients. Another possible effect of mutation is an increased spreading speed. Taking again a system with only two morphs (as in Deforet et al. (2019), but with a mutation term), typically an R-specialist and a *D*-specialist, Elliott and Cornell (2012) and Morris et al. (2019) investigated the effect of varying *R* and *D* on the spreading speed. They found that the system would spread faster in the presence of both phenotypes than just one phenotype would spread in the absence of mutation for certain combination of *R* and *D* values. In a similar way, using the results of Girardin (2017), Keenan and Cornell (2021) considered the *R — D* trades-off in the case of *N* phenotypes in a homogeneous environment, and obtained some conditions on the curvature of the trade-off curve (*D,R*(*D*)) such that this “anomalous” faster speed emerges. Here, although the trade-off curve (*D, R*(*D*)) has positive curvature, we did not observe this phenomenon: in all of our simulations of Figures 3–5, the speed is reduced when *μ* > 0, compared to the speed of the fastest trait when *μ* = 0. The theoretical results of Keenan and Cornell (2021) require a vanishing small mutation rate, and their numerical results use a mutation coefficient 10^-6^ (to be compared with *μ*/(*δ_x_*)^2^ = (0.1)/ (0.1)^2^ = 10 in our continuous framework), which may explain these differences.

Future works could consider a more detailed analysis of the lineages that pull the forefront, to determine for instance if the *D*-specialists in Figures 3 and 5 are produced by mutation from *R*-specialists or correspond to a self-sustaining fraction of the population. In that respect, one could reconstruct the genealogies of the fractions composing the population using the methods in (Roques et al., 2012). We recall that our results depend on the assumptions about the form of the dispersal and growth rate functions and the fragmentation definition. For instance, we do not take an Allee effect into account. It is demonstrated to have important consequences on the invasions dynamics and especially on the lineages that compose the forefront (Andrade-Restrepo et al., 2019; Chuang and Peterson, 2016; Roques et al., 2012).

## 7 Data script and code availability

Data and script are available online: DOI 10.17605/OSF.IO/V6N4M, the webpage hosting the data: https://osf.io/v6n4m/. Supplementary information is available online: https://www.biorxiv.org/content/10.1101/2022.04.07.487471v2.supplementary-material.

