## Supplementary information for "Spatial heterogeneity alters the trade-off between growth and dispersal during a range expansion"

#### Appendix A: Effect of the amplitude on the fragmentation

Here, we investigate the role of the amplitude of fragmentation on influencing the fastest phenotype leading the forefront. Now, the spatial fragmentation in environmental conditions impacting the consumer growth rate $R$ and its mobility $D$ is slightly modified introducing the variable $a \in [0-1]$, which scales up the amplitude of the spatial fragmentation. Thus, simulation scenarios are redefined as follows:

- Scenario $H$: spatially homogeneous coefficients. In this case,

$$R(x,y) = R_h(y) = R_0 + R_g(y) \text{ and } D(x,y) = D_h(y) = D_0 + D_g(y).$$

- Scenario $R_{het}$: fragmented growth and homogeneous dispersal. In this case,

$$R(x,y) = R_0 + R_g(y) + a\, R_s(x/L) \text{ and } D(x,y) = D_h(y) = D_0 + D_g(y).$$

- Scenario $D_{het}$: homogeneous growth and fragmented dispersal. In this case,

$$R(x,y) = R_h(y) = R_0 + R_g(y) \text{ and } D(x,y) = D_0 + D_g(y) + a\,D_s(x/L).$$

We numerically simulate the model of Equation (1) with the growth rate $R(x,y)$ and dispersal $D(x,y)$ functions to assess how the fastest phenotype leading the forefront is affected by the shape of the environmental fragmentation (period $L$ and amplitude $a$) and the trade-off strength (distance among the optimum $d$). Results are presented in Figure A1 for the different scenarios.

In the scenario $R_{het}$ (Figure A1 panels A and B), for $L = 2$, the fragmentation amplitude $a$ does not produce any effect on the dynamics: when $d < d_{cr}$, the $R - D$ trade-off is always in favor of the generalist strategy $y* = 0$; when $d > d_{cr}$ the fastest phenotype corresponds to $y* = O_R$ (Figure A1, panel A). Instead, for $L = 10$, when $d > d_{cr}$, the fastest phenotype could be either $y* = O_R$ or $y* = O_D$ (Figure A1, panel B). Then, the behavior depends on the values of the amplitude of fragmentation ($a$): low values favor a strategy where $y* = O_R$ while high values favor a strategy where $y* = O_D$.

In the scenario $D_{het}$ (Figure A1, panels C and D), for $d < d_{cr}$, the fastest phenotype is always $y* = 0$. For $d > d_{cr}$, as before, the fastest phenotype could be either $y* = O_R$ or $y* = O_D$ depending on the value of $a$.

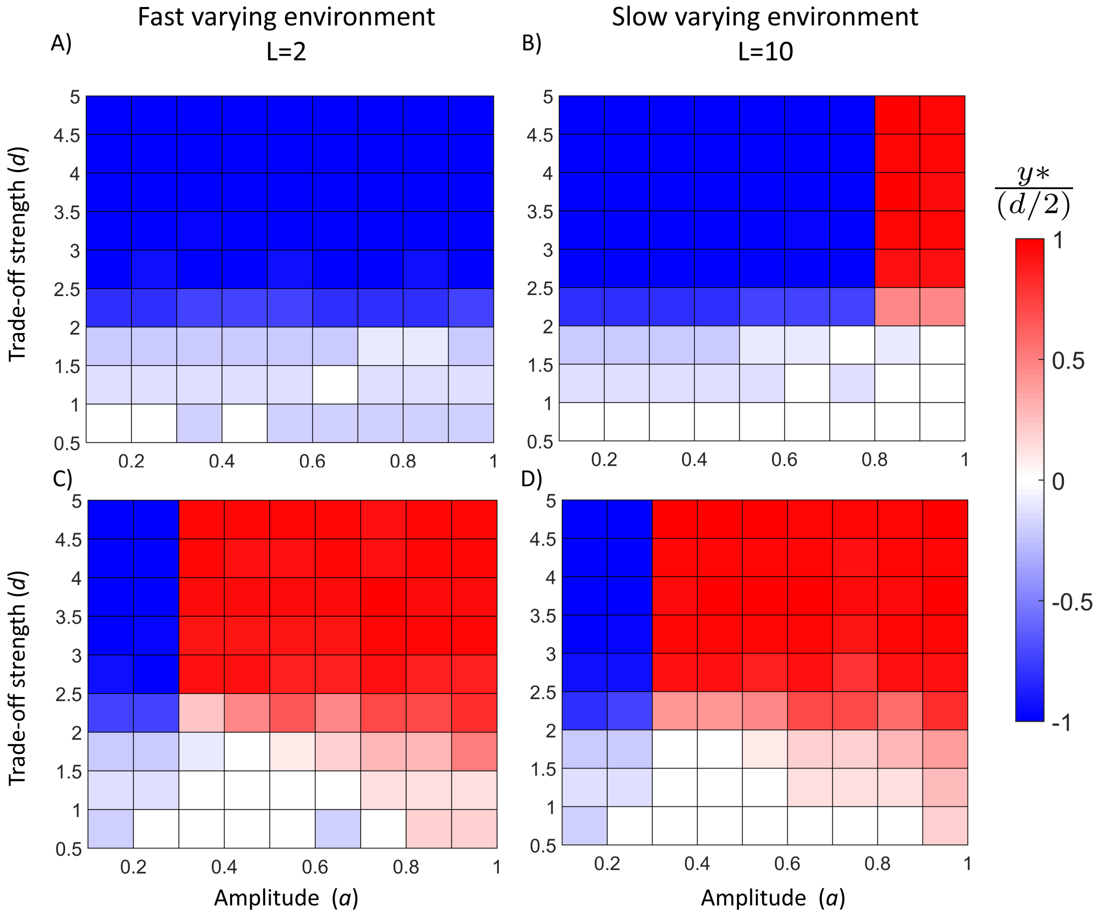


Figure A1: **Effect of the amplitude of heterogeneity $a$ without mutation ($\mu = 0$).** Representation of the fastest phenotype leading the forefront ($y* \to O_R$, blue shades; $y* \to O_D$, red shades; $y* \to 0$ white) as a function of the amplitude of fragmentation ($a = [0.1 - 1]$) and the trade-off strength (distance among the optimum values, $d = [0.5 - 5]$) without mutation ($\mu = 0$). We report scenario $R_{het}$ (panel A and B) and $D_{het}$ (panel C and D) for $L = 2$ (panel A and C) and $L = 10$ (panel B and D). These results were obtained through a numerical simulation of Equation (1).

{fig:a-d}

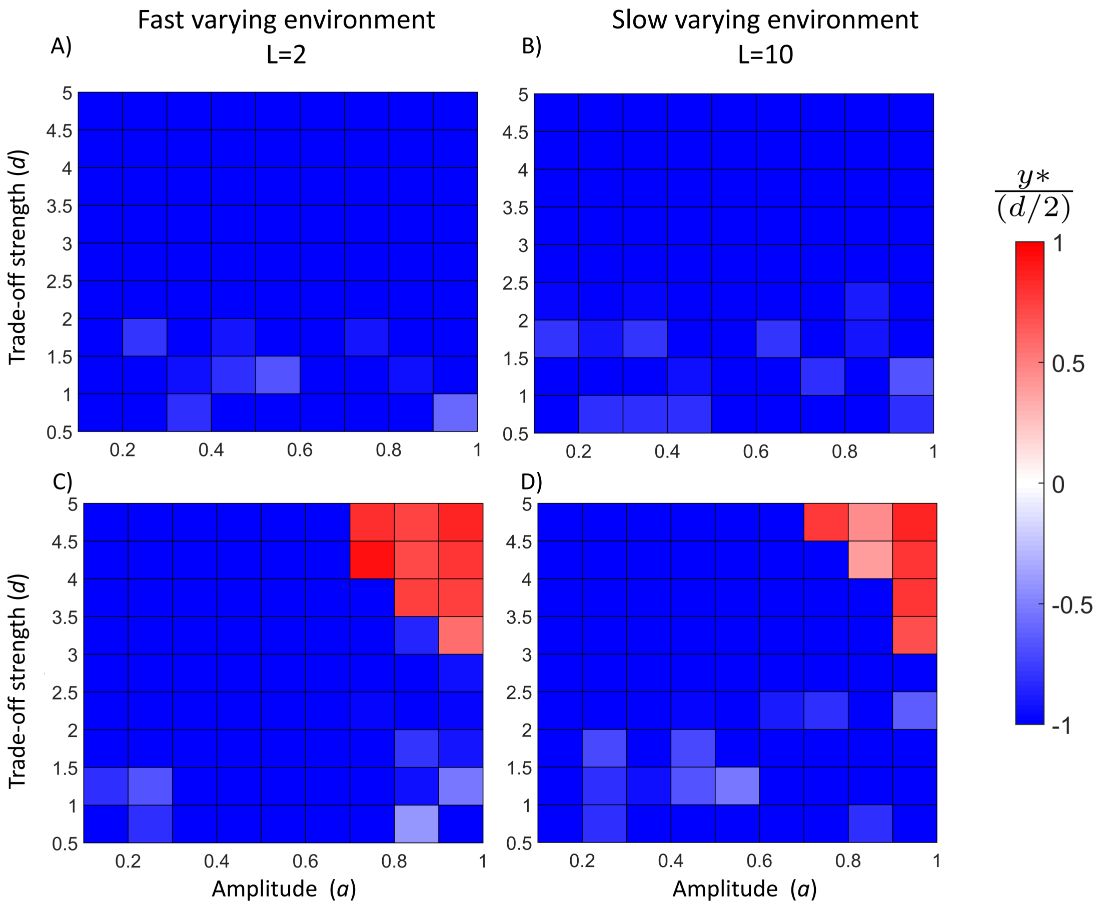


Figure A2: **Effect of the amplitude of heterogeneity $a$ without mutation ($\mu > 0$).** Representation of the fastest phenotype leading the forefront ($y* \to O_R$, blue shades; $y* \to O_D$, red shades; $y* \to 0$ white) as a function of the amplitude of fragmentation ($a = [0.1 - 1]$) and the trade-off strength (distance among the optimum values, $d = [0.5 - 5]$) with mutation ($\mu = 0.1$). We report scenario $R_{het}$ (panel A and B) and $D_{het}$ (panel C and D) for $L = 2$ (panel A and C) and $L = 10$ (panel B and D). These results were obtained through a numerical simulation of Equation (1).

{fig:a-d_mu}

### Appendix B: Spreading speed

*Scenario H: Homogeneous case*

In the absence of mutation ($\mu = 0$), it is known that the spreading speed of the solution of (6) in the Homogeneous case (*i.e.* $R = R_h(y)$ and $D = D_h(y)$) associated with a phenotype $y$ is $v(y) = 2\sqrt{R_h(y)\, D_h(y)}$ (Kolmogorov et al., 1937). Depending on $d$ and $\sigma$ (see Equations (2)-(3)), there may be only one fastest phenotype $y^*$ in 0 or two optima at $\pm d/2$.

Note that $D_h(y) = R_h(-y)$, thus:

$$(R_h(y)\, D_h(y))' = R_h'(y)R_h(-y) - R_h(y)R_h'(-y).$$

For $y = 0$, this quantity is equal to 0. Computing the second derivative, *i.e.*,

$$(R_h\, D_h)''(y) = R_h''(y)R_h(-y) - 2\, R_h'(y)R_h'(-y) + R_h''(-y)R_h(y),$$

at $y = 0$, yields $(R_h\, D_h)''(0) = 2(R_h''(0)R_h(0) - R_h'(0)^2)$. Thus, we observe that:

$$\begin{cases} (R_h\, D_h)''(0) > 0, & \text{if } d > 2\sigma\sqrt{1 + 2\, W\left(\frac{1}{2\, R_0} e^{-1/2}\right)}, \\ \\ (R_h\, D_h)''(0) < 0, & \text{if } d < 2\sigma\sqrt{1 + 2\, W\left(\frac{1}{2\, R_0} e^{-1/2}\right)}, \end{cases} \tag{B1}$$

with $W$ the principal branch of the Lambert function. Thus, there exists a threshold on the optimum distance given by $d_{cr} = 2\sigma\sqrt{1 + 2\, W\left(\frac{1}{2\, R_0} e^{-1/2}\right)}$. When $d < d_{cr}$, 0 is a (local) maximum of the speed, while if $d > d_{cr}$, we conclude that the speed has two maxima, which are symmetric with respect to 0. As $(R_h\, D_h)'(d/2) < 0$, these maxima are reached in the region $(-d/2, d/2)$.

*Scenario $R_{het}$: Fragmented R, homogeneous D*

In the local Equation (6), when the coefficient $R(x,y)$ is spatially fragmented and $L-$periodic, we adopt the formula given for the spreading speed in Berestycki and Hamel (2005). To state this formula, we first have to define a differential operator $\mathcal{L}_\lambda$, acting on functions $\phi(x,y)$ which are $L-$periodic in $x$ and satisfy no-flux boundary conditions at the boundaries $y = y_{\min}, y_{\max}$. For any $\lambda > 0$, this operator is defined by:

$$\mathcal{L}_\lambda(\phi) := D(y)\,\partial_{xx}\phi + \mu\,\partial_{yy}\phi + 2\,\lambda\,D(y)\,\partial_x\phi + [\lambda^2 D(y) + R(x,y)]\phi. \tag{B2}$$

The spreading speed can then be computed by the Gärtner-Freidlin formula:

$$V = \min_{\lambda>0} \frac{k(\lambda)}{\lambda}, \tag{B3}$$

with $k(\lambda)$ the principal eigenvalue (the unique eigenvalue associated with a positive eigenfunction) of $\mathcal{L}_\lambda$.

When $\mu = 0$, more tractable theoretical formulas can be obtained to determine the spreading speed $v(y)$ associated with a given phenotype $y$ at the limit of rapidly varying and slowly varying environments (*i.e.* $L \to 0$ and $L \to \infty$, respectively).

1. **Rapidly varying environment** $L \to 0$

   We refer to Smaily et al. (2009) work, where the spreading speed $v_0(y)$ of each phenotype is derived analytically:

   $$v_0(y) = 2\sqrt{D_h(y)\,\overline{R}_x(y)}, \tag{B4}$$

   with $\overline{R}_x(y)$ the mean value of the growth rate, averaged over space:

   $$\overline{R}_x(y) = R_0 + R_g(y) + \int_0^1 R_s(x)\,dx = R_h(y),$$

as $R_s(x)$ has mean value 0. Finally, the speed is the same as in the homogeneous case (that is, with $R_s \equiv 0$):

$$v_0(y) = 2\sqrt{R_h(y)\, D_h(y)}.$$

2. **Slowly varying environment** $L \to +\infty$

   An explicit formula for the limit spreading speed $v_\infty(y)$ of each phenotype as $L \to +\infty$ can be derived analytically using the formulation of Hamel et al. (2010). It is given by the expression:

$$v_\infty(y) = 4\sqrt{D_h(y)} \times \frac{(R^+(y))^2 + (R^-(y))^2 + (R^+(y) + R^-(y))\,\sqrt{\triangle(y)}}{(R^+(y) + R^-(y) + 2\sqrt{\triangle(y)})^{\frac{3}{2}}}$$

   with $R^+(y) = R_h(y) + R_0$, $R^-(y) = R_h(y) - R_0$ and $\triangle(y) = (R^+(y))^2 + (R^-(y))^2 - R^+(y)\,R^-(y)$.

*Scenario $D_{het}$: Homogeneous R, fragmented D*

The Fokker-Planck diffusion term in the local Equation (6), can be rewritten as:

$$\partial_{xx}(D(x,y)\,c(t,x,y)) = \partial_x(D(x,y)\,\partial_x c(t,x,y)) + \partial_x(c(t,x,y)\,\partial_x D(x,y)). \qquad \text{(B5)}$$

The term $\partial_x(D(x,y)\,\partial_x c(t,x,y))$ corresponds to Fickian diffusion operator. Though both types of diffusion terms are found in the ecological literature, Fokker-Planck diffusion $\partial_{xx}(D(x,y)\,c(t,x,y))$ naturally emerges from Brownian motion with space-dependent mobility, and seems better adapted to describe individual movements (see Roques, 2013; Turchin, 1998), whereas Fickian diffusion emerges from flux considerations and seems more adapted to describe heat conduction in fragmented media. The main difference between these two operators is that Fickian diffusion tends to homogenize the density

$c(t,x,y)$ (with respect to the diffusion variable $x$). The additional term

$$\partial_x(c(t,x,y)\,\partial_x D(x,y))$$

in (B5) corresponds to a spatially-periodic transport term which is oriented towards the lower values of $D$. As the standard formula for the spreading speed with fragmented diffusion in (Berestycki and Hamel, 2005) and the formulas in the limiting cases (Hamel et al., 2011; Smaily et al., 2009) are only available for Fickian diffusion operators, we used these formulas in Table 1 (scenario $D_{het}$), thereby neglecting effect of the transport term $\partial_x(c(t,x,y)\,\partial_x D(x,y))$ on the spreading speeds $v(y)$. The corresponding equation, with mutation rate $\mu = 0$ is:

$$\partial_t c(t,x,y) = \partial_x(D(x,y)\,\partial_x c(t,x,y)) + c(t,x,y)\,(R(x,y) - \gamma\, c(t,x,y))\,. \tag{B6}$$

1. **Rapidly varying environment** $L \to 0$

   The spreading speed $v_0(y)$ of each phenotype can be derived from Smaily et al. (2009):

   $$v_0(y) = 2\sqrt{R_h(y)\,\langle D_1\rangle_H(y)},$$

   with $\langle D_1\rangle_H(y) = \left(\int_0^1 \frac{1}{D_1(x,y)}\,dx\right)^{-1}$ the harmonic mean of $x \mapsto D_1(x,y) = D_0 + D_g(y) + D_s(x)$.

2. **Slowly varying environment** $L \to +\infty$

   In this case, we use the results in Hamel et al. (2011), which show that:

   $$v_\infty(y) = 2\sqrt{R_h(y)}\,\langle\sqrt{D_1}\rangle_H(y),$$

   with $\langle\sqrt{D_1}\rangle_H(y) = \left(\int_0^1 \frac{dx}{\sqrt{D_1(x,y)}}\right)^{-1}$ the harmonic mean of $x \mapsto \sqrt{D_1(x,y)} = \sqrt{D_0 + D_g(y) + D_s(x)}$.

### Appendix C: Long simulation for checking phenotype composition behind the front

Here, we check the phenotype selection strategy behind the front to confirm the selection of R-strategy under the scenario $D_{het}$ for a slowly varying environment. We simulated the scenario $D_{het}$ without mutation ($\mu = 0$) for a longer time (we took twice as long as the time used in the simulations in the main text, $T = 120$) in a bigger domain (we used double the space compared to the spatial domain used in the simulations in the main text, $x \in [0, 500]$). The results presented in figure C1 is close to the ones presented in the main text: on the forefront, the selected strategy is the dispersal one, but at the very back of the front phenotypes select R-strategies as $y* < 0$.

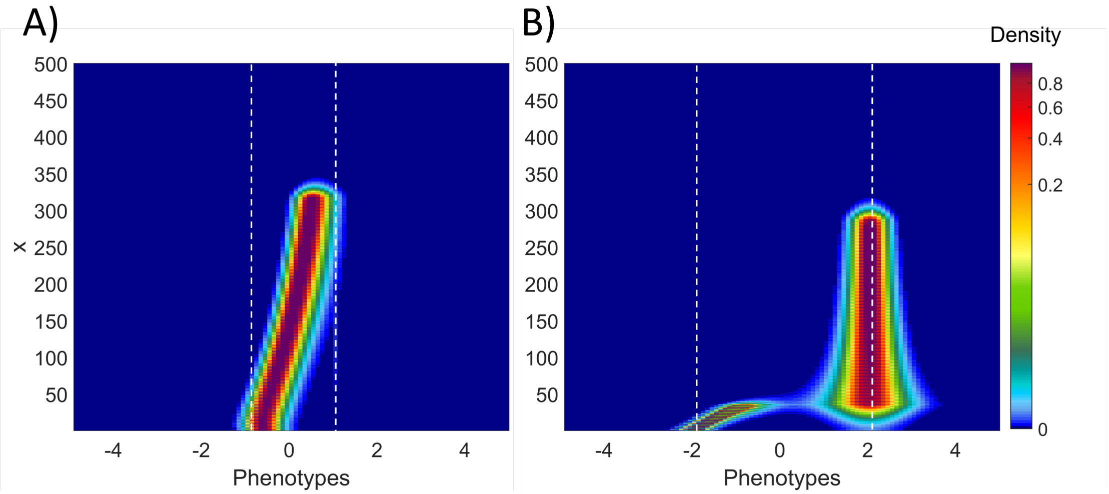


Figure C1: **Population density under the scenario $D_{het}$ in a rapidly varying environment in a bigger spatial space and for longer time without mutation.** We report scenario $D_{het}$ for $L = 2$ considering $d = 2$ in panel A and $d = 4$ in panel B at time $T_{sim} = 60$ in a spatial domain equal to $x \in [0, 500]$ for $\mu = 0$. White dashed lines highlight the corresponding optimum trait values (*i.e*, $O_D = d/2$ and $O_R = -d/2$). These results were obtained through a numerical simulation of Equation (1).

{fig:long}
